# Replacing the Calvin cycle with the reductive glycine pathway in *Cupriavidus necator*

**DOI:** 10.1101/2020.03.11.987487

**Authors:** Nico J. Claassens, Guillermo Bordanaba-Florit, Charles A. R. Cotton, Alberto De Maria, Max Finger-Bou, Lukas Friedeheim, Natalia Giner-Laguarda, Martí Munar-Palmer, William Newell, Giovanni Scarinci, Jari Verbunt, Stijn T. de Vries, Suzan Yilmaz, Arren Bar-Even

## Abstract

Formate can be directly produced from CO_2_ and renewable electricity, making it a promising microbial feedstock for sustainable bioproduction. *Cupriavidus necator* is one of the few biotechnologically-relevant hosts that can grow on formate, but it uses the inefficient Calvin cycle. Here, we redesign *C. necator* metabolism for formate assimilation via the highly efficient synthetic reductive glycine pathway. First, we demonstrate that the upper pathway segment supports glycine biosynthesis from formate. Next, we explore the endogenous route for glycine assimilation and discover a wasteful oxidation-dependent pathway. By integrating glycine biosynthesis and assimilation we are able to replace *C. necator*’s Calvin cycle with the synthetic pathway and achieve formatotrophic growth. We then engineer more efficient glycine metabolism and use short-term evolution to optimize pathway activity, doubling the growth yield on formate and quadrupling the growth rate. This study thus paves the way towards an ideal microbial platform for realizing the formate bioeconomy.

## Introduction

Microbial biosynthesis offers an environmentally friendly alternative to fossil-based production. However, the limited availability and questionable sustainability of microbial feedstocks hamper the expansion of biotechnological production and the establishment of a circular carbon economy. The common substrates for microbial bioproduction are plant-based sugars, the utilization of which competes with food supply and necessitates vast land use that negatively impacts the environment. Moreover, alternative feedstocks, such as lignocellulosic biomass, suffer from crucial drawbacks, such as difficult and expensive processing^1^. A fundamental limitation of all photosynthesis-based resources is the low energy conversion efficiency associated with this process, typically below 1%^2,3^.

Electromicrobial production has gained attention as an alternative route towards sustainable biotechnology^4,5^. This strategy is based on the use of two key feedstocks: CO_2_-free electricity – e.g. from solar, wind, hydro – the production of which is rapidly growing, and CO_2_, a virtually unlimited carbon source, captured either from point sources or directly from air. Some microbes can grow by receiving electrons directly from a cathode; however, low current densities limit the economic viability of this approach^6,7^. A more feasible option is the electrochemical production of small reduced compounds^6^ that are subsequently fed to microbes and then converted into value-added chemicals. Among the possible mediator compounds, hydrogen, carbon monoxide, and formate can be produced at high efficiency and rate^8^. Whereas hydrogen and carbon monoxide are gases of low solubility, formate is completely miscible and can be readily introduced to microbial cells without mass transfer limitations and without major safety concerns^9^. Hence, establishing a “formate bio-economy” has been proposed as a route towards realizing a circular carbon economy^10^.

*Cupriavidus necator* (formerly *Ralstonia eutropha*) is one of the very few formatotrophic microorganisms that have been extensively explored for biotechnological use^11,12^. *C. necator* has been industrially utilized for the production of polyhydroxybutyrate and further engineered for the biosynthesis of various commodity and specialty chemicals such as isopropanol or terpenoids^13–15^. However, conversion of formate to biomass and products by this bacterium is hampered by low yields due to the use of the inefficient and ATP-wasteful Calvin cycle (Fig. 1)^16,17^.

**Figure 1.**
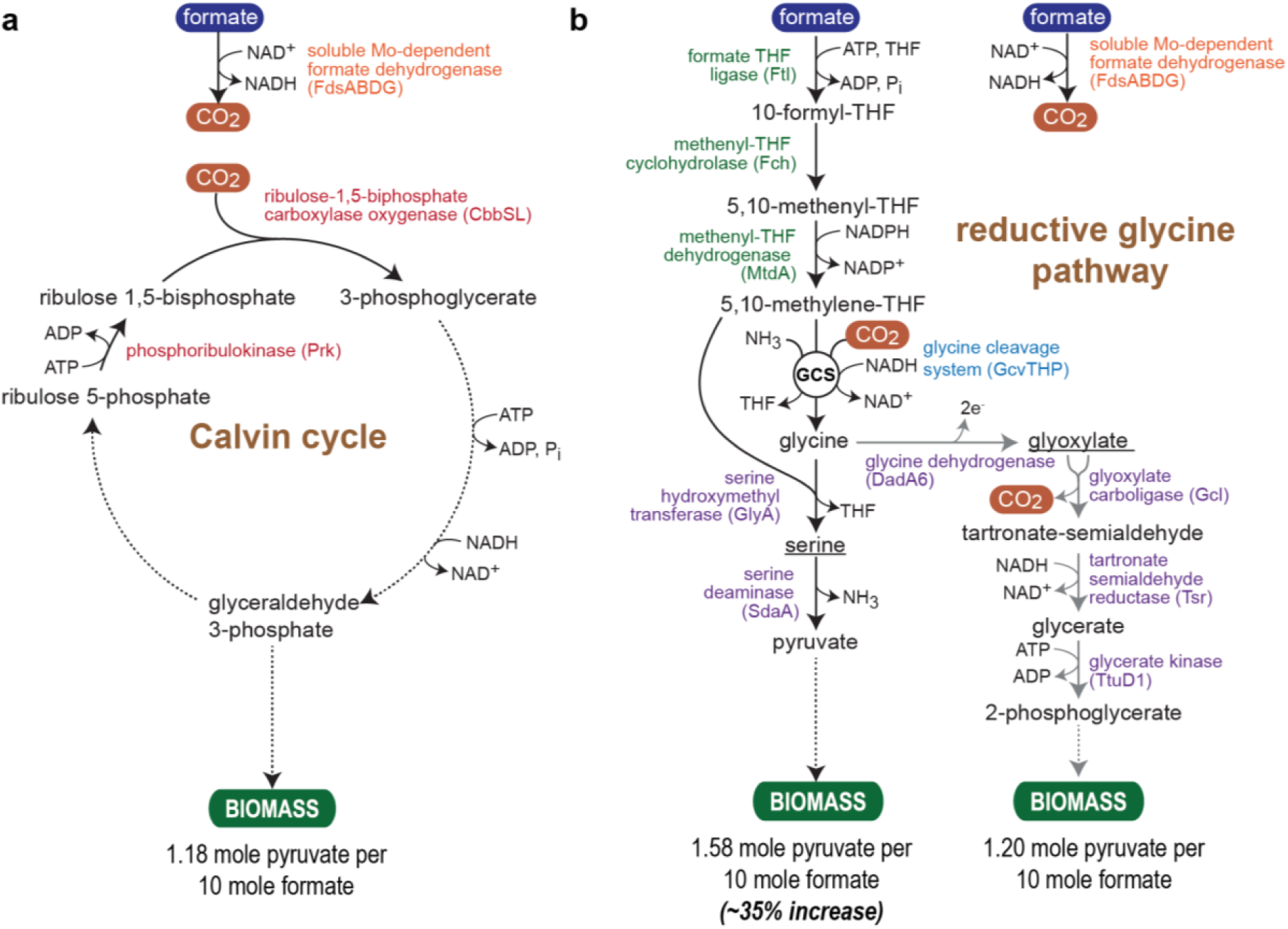
Structures of the native Calvin cycle and synthetic reductive glycine pathway (rGlyP). (**a**) Simplified diagram showing formate dehydrogenase and the Calvin cycle active in WT *C. necator* (**b**) Two variants of the rGlyP are shown: (i) the original design in which glycine is assimilated via serine and pyruvate and (ii) an alternative variant where glycine is first oxidized to glyoxylate, which is then assimilated via the glycerate pathway. *C. necator* protein names are written in brackets, except for the enzymes catalyzing formate metabolism to methylene-THF (green), where the proteins are from *M. extorquens.* Pyruvate molar yields per formate are theoretical estimates based on the number of formate molecules directly assimilated (Calvin: 0, rGlyP-ser: 2, rGlyP-glyox: 2), used for NADH production (Calvin: 5, rGlyP-ser: 1, rGlyP-glyox: 3), used for NADPH production (Calvin: 0, rGlyP-ser: 2*(1.33), rGlyP-glyox: 2*(1.33), where NADPH requires a proton translocation) and used for ATP production (Calvin: 7/2, rGlyP-ser: 2/2, rGlyP-glyox: 2/2, where a P/O ratio of 2 was assumed). Abbreviation: GCS, Glycine cleavage system; THF, tetrahydrofolate.

We previously proposed the synthetic reductive glycine pathway (rGlyP) as the most energy-efficient formate assimilation route that can operate under aerobic conditions^16^. In this pathway, formate is condensed with tetrahydrofolate (THF) and further reduced to methylene-THF. Then, the glycine cleavage system (GCS), operating in the reductive direction, condenses methylene-THF with CO_2_ and ammonia to produce glycine. Glycine is subsequently metabolized to serine and deaminated to pyruvate which serves as a biomass precursor (Fig. 1). The rGlyP is considerably more efficient than the Calvin cycle: while the latter pathway can maximally generate ∼1.2 mole pyruvate per 10 mole formate consumed, the rGlyP can theoretically produce ∼1.6 mole pyruvate for the same amount of feedstock, i.e., increasing yield by ∼35% (Fig. 1).

Here, we replace the Calvin cycle of *C. necator* with the rGlyP. First, we show that overexpression of the upper segment of the pathway enables an otherwise glycine-auxotrophic *C. necator* to utilize formate for glycine biosynthesis. We then evolve *C. necator* to utilize glycine as a carbon source, revealing that, rather than being converted to serine and pyruvate, this amino acid is first oxidized to glyoxylate, which is subsequently assimilated via the well-known glycerate route^18,19^ (Fig. 1). Next, we construct a strain in which the Calvin cycle was disrupted and is thus unable to grow on formate. Integration of the two segments of the rGlyP restores the formatotrophic growth of this strain, albeit at a low growth rate. We further optimize pathway activity by shifting overexpression from a plasmid to the genome, forcing glycine assimilation via serine, and conducting a short-term adaptive evolution. Our final strain displays a biomass yield on formate equivalent to that of the WT strain using the Calvin cycle, hence confirming the recovery of the growth phenotype after the fundamental rewiring of cellular metabolism towards the use of the synthetic route. Our study therefore paves the way towards a highly efficient platform strain for the production of value-added chemicals from CO_2_.

## Results

### Engineering conversion of formate to glycine

First, we aimed to establish the upper segment of the rGlyP which converts formate into glycine (Fig. 1). *C. necator* natively harbors all the enzymes of this segment with the exception of formate-THF ligase, which needs to be heterologously expressed. In a previous study in *E. coli* we observed that the native FolD enzyme – encoding for a bifunctional methylene-THF dehydrogenase / methenyl-THF cyclohydrolase – is not suitable for carrying high flux in the reductive direction^20^. Hence, we decided to heterologously express three enzymes from *Methylobacterium extorquens* that naturally support high reductive flux from formate to methylene-THF (Fig. 1): Ftl (formate–THF ligase), Fch (methenyl-THF cyclohydrolase), and MtdA (methylene-THF dehydrogenase)^21^. The genes encoding for these enzymes were cloned from *M. extorquens* and assembled into a synthetic operon, in a plasmid named pC1, using four different constitutive promoters of varying strength (Methods and Fig. S1, S2). For the subsequent conversion of methylene-THF to glycine, we overexpressed the native genes of the GCS (*gcvT, gcvH*, and *gcvP*) from a synthetic operon on a second plasmid (termed pC2), which was constructed with three constitutive promoters of different strength (Methods and Fig. S1, S2).

We then generated a *C. necator* strain auxotrophic to glycine by deleting the genes encoding for serine hydroxymethyltransferase (*ΔglyA*), threonine aldolase (*ΔltaA*), and glycine acetyltransferase (*Δkbl*) (Fig. 2a). This strain could grow only when supplemented with glycine (Fig. 2b, brown line vs. grey line). We transformed this strain with pC1 and pC2 carrying different combinations of promoters. All strains harboring pC2 with promoter p_3_ (the weakest promoter tested), regardless of the promoter in pC1, were able to grow with fructose as main carbon source and formate and CO_2_ as glycine source (at 10% CO_2_ and 100 mM bicarbonate; high CO_2_ concentration is required to thermodynamically and kinetically push the glycine cleavage system in the reductive direction). Yet, the strain harboring the weak promoter p_14_ in pC1 showed the best growth. The growth of the strain carrying pC1-p_3_ and pC2-p_14_ on fructose and formate but in the absence of glycine (green line in Fig. 2b), was almost identical to the positive control, in which glycine was added to the medium (brown line in Fig. 2b). Growth was also possible in the absence of fructose, in which case formate served both as a carbon source for glycine biosynthesis and as a source of reducing power source to support growth via the Calvin cycle (orange line in Fig. 2b). Interestingly, transformation of the glycine auxotroph strain with pC1-p_3_ alone sufficed to support glycine biosynthesis from formate, albeit at a low rate (blue line in Fig. 2b), indicating that the native expression of the GCS supports at least some reductive activity.

**Figure 2.**
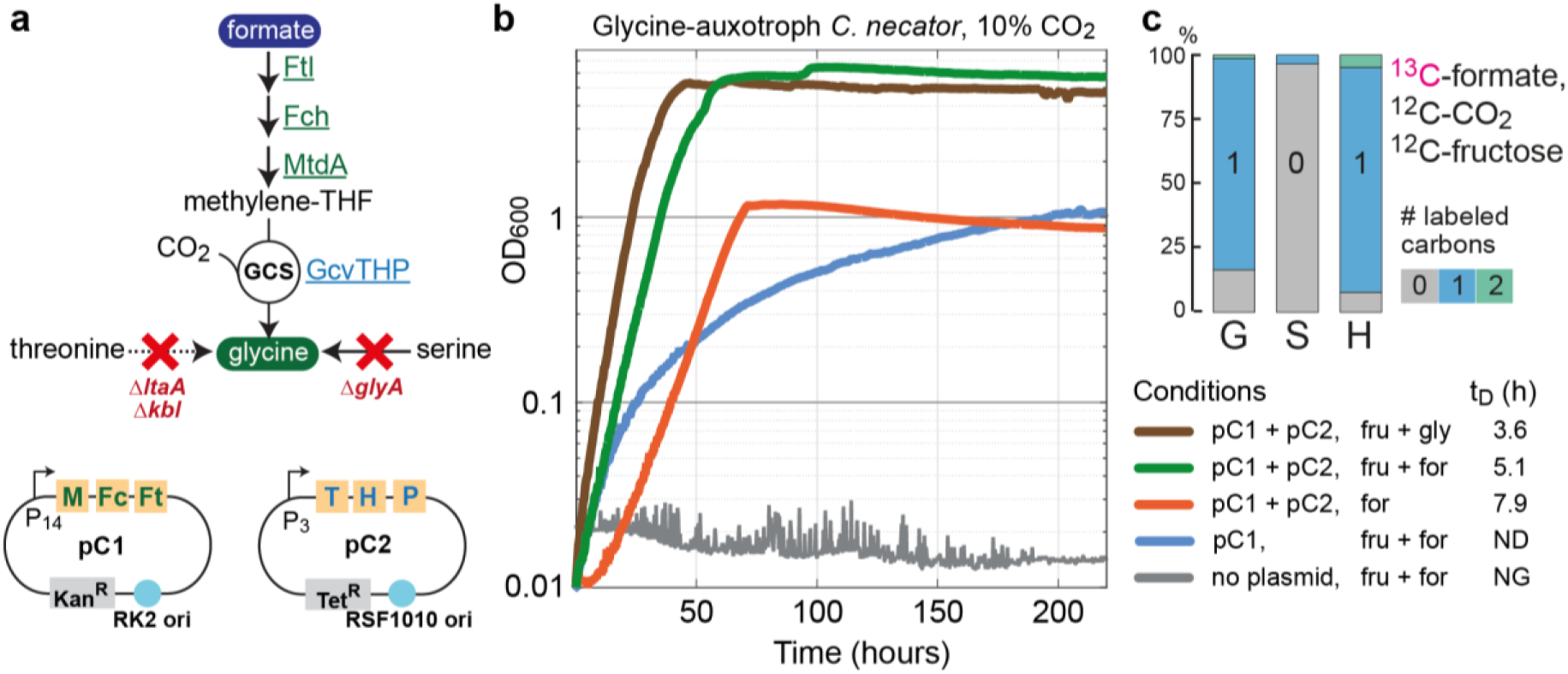
Engineering glycine biosynthesis from formate. (**a**) Selection scheme for the selection of glycine biosynthesis from formate in an otherwise glycine auxotroph *C. necator* strain (deleted in *glyA, ltaA*, and *kbl*). Proteins that were overexpressed are underlined. The plasmids resulting in the fastest growth (pC1-p_14_, pC2-p_3_) are shown below (‘M’ corresponds to *mtdA*; ‘Fc’ to *fch*; ‘Ft’ to *ftl*; ‘T’ to *gcvT*; ‘H’ to *gcvH*; and ‘P’ to *gcvP*). (**b**) Overexpression of the pathway enzymes rescue the growth of a glycine auxotroph with formate as glycine source. Growth experiments were conducted in 96-well plate readers in minimal medium (JMM) supplemented with 100 mM bicarbonate and 10% CO_2_ in the headspace. When added, the concentrations of the different carbon sources were 20 mM fructose (‘fru’), 8 mM glycine (‘gly’), and 80 mM formate (‘for’). ‘ND’ corresponds to ‘not determined’, and ‘NG’ to’ no growth’. Growth experiments were performed in triplicates and showed identical growth curves (±5%); hence, representative curves and average doubling times (t_D_) are shown. All experiments were repeated independently at least three times and showed highly similar growth behavior. (**c**) ^13^C-labeling of proteinogenic amino acids after cultivating the glycine auxotroph harboring pC1 and pC2 on unlabeled fructose and CO_2_ as well as ^13^C-formate. The labeling pattern confirms the operation of the upper segment of the pathway, as glycine (G) and histidine (H, harboring a carbon derived from formyl-THF) are dominantly labeled once, while serine (S) is non-labeled. We note that the small fraction of non-labeled glycine might be attributed to the amination of glyoxylate via the promiscuous activity of aminotransferase enzymes; we did not delete isocitrate lyase, the source of cellular glyoxylate, as we found this deletion to hamper cell viability. Labeling experiments are averages from duplicates.

To confirm that glycine as well as the cellular C_1_ moieties are indeed generated from formate assimilation, we conducted a ^13^C-labeling experiment. We cultivated the strain harboring pC1-p_3_ and pC2-p_14_ on unlabeled fructose and CO_2_ as well as ^13^C-formate. We measured the labeling pattern of glycine and histidine (the latter contains a carbon derived from 10-formyl-THF) as well as serine, which was expected not to be labeled. We found glycine and histidine to be almost completely once labeled and serine to be unlabeled, thus confirming that formate is assimilated to the THF pool and into glycine (Fig. 2c).

### Exploring growth on glycine

Next, we turned our attention to the downstream segment of the rGlyP, that is, assimilation of glycine into biomass. We explored the native capacity of *C. necator* to grow on glycine. We inoculated twelve parallel cultures of *C. necator* on a minimal medium with glycine as a sole carbon source. Initially, no growth was observed. However, after a week, three of the inoculated cultures started growing. When reinoculated into a fresh medium with glycine, these cultures started growing immediately, possibly due to genetic adaptation. We performed whole-genome sequencing of these three strains and found that each had a sense mutation in either *gltR1* or *gltS1* (Supplementary Data 1). These two genes reside in the same operon and encode for a dual-component signal-regulator system that activates the expression of *gltP1*, encoding for a putative dicarboxylate/amino acid transporter (Fig. 3a). While we detected several other mutations in specific strains (Supplementary Data 1), they did not appear in all three strains and hence they probably have lower contribution to growth on glycine.

**Figure 3.**
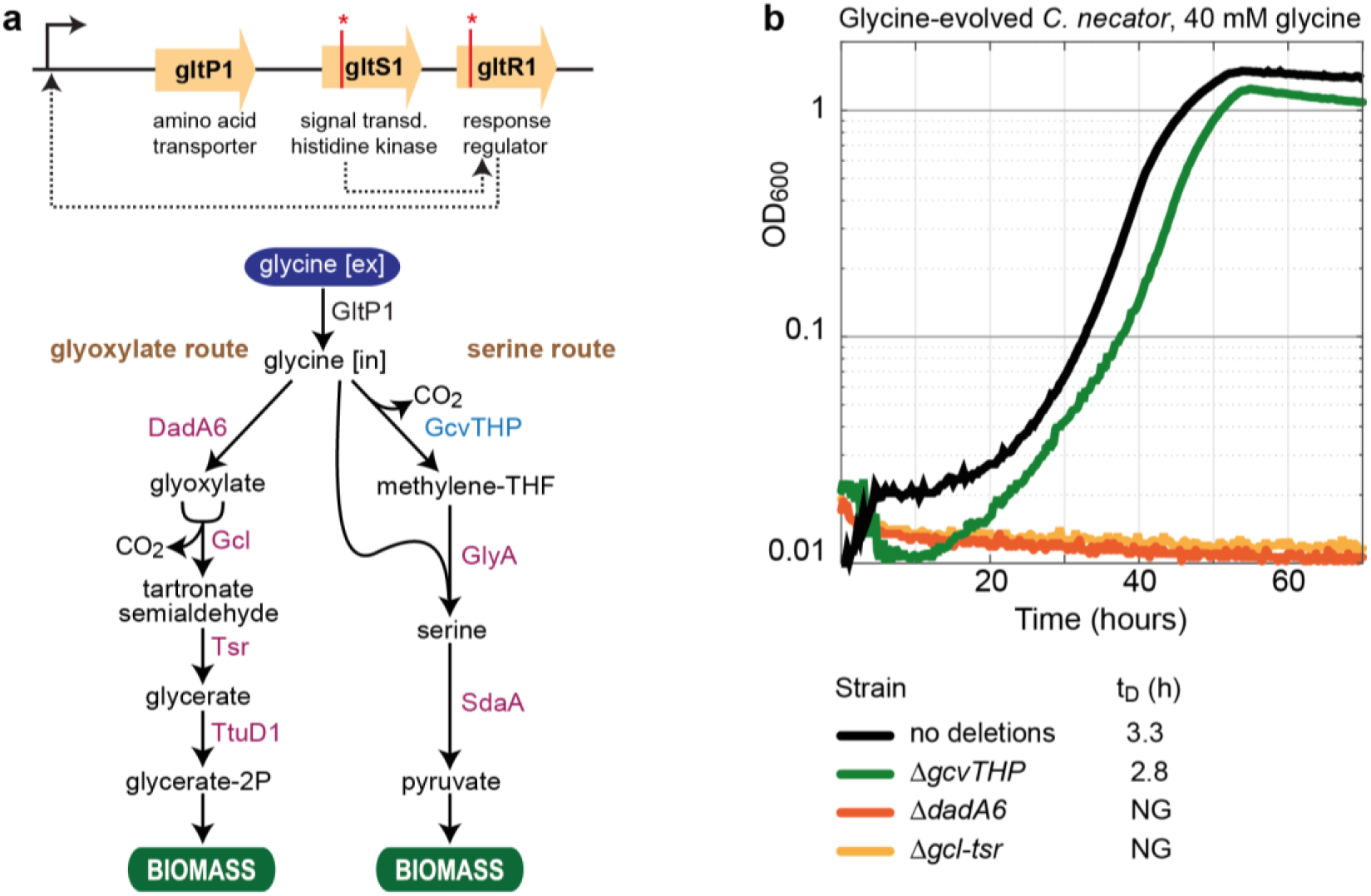
*C. necator* can grow on glycine after short evolution via an oxidative pathway. (**a**) Three independently evolved strains harbored mutations either in *gtlR1* or *gtlS1* (Supplementary Data 1), the regulator system of the amino acid transporter *gtlP1*. Within the cell, glycine can be oxidized to glyoxylate by DadA6 and assimilated through the glycerate pathway, the transcription of which was highly upregulated (Supplementary Data 2). Alternatively, glycine can be metabolized via glycine cleavage (by the GCS) and converted to serine and pyruvate. (**b**) Growth of the evolved strain on glycine with or without further gene deletions, as indicated in the legend. This experiment confirms that glycine is assimilated via the “glyoxylate route” rather than the “serine route”. Growth experiments were conducted in 96-well plate readers on a minimal medium (JMM) supplemented with 40 mM glycine and 10% CO_2_ in the headspace. Growth experiments were performed in triplicates and showed identical growth curves (±5%); hence, representative curves and average doubling times (t_D_) are shown. All experiments were repeated independently at least three times and showed highly similar growth behavior.

We analyzed the transcriptome of one of the evolved strains. In comparison to the WT strain growing on pyruvate, we observed >1,000-fold increase of *gltP1* transcription in the evolved strain growing on glycine. It therefore seems that GltP1 acts as a glycine transporter and that glycine, once within the cell, activates a native route for its metabolism. To uncover this endogenous glycine assimilation pathway we checked which other genes were significantly overexpressed in the glycine-assimilating strain (Supplementary Data 2). *dadA6*, encoding for a putative FAD-dependent D-amino acid dehydrogenase, was amongst the 10 most upregulated genes (∼200-fold increase in transcript abundance). Given that a similar enzyme from *Bacillus subtilis* was demonstrated to oxidize both D-amino acids and glycine^22^, we speculated that DadA6 can also oxidize glycine to glyoxylate. In addition, the genes encoding for glyoxylate carboligase (*gcl*), tartronate semialdehyde reductase (*tsr*), and glycerate kinase (*ttuD1*) were amongst the 10 most upregulated genes (>1600-fold, ∼400-fold, ∼300-fold increased transcript, respectively). This led us to speculate that glycine is assimilated via oxidation to glyoxylate, followed by the activity of the well-known glycerate pathway^18,19^ (Fig. 3a).

To test whether DadA6 can indeed catalyze glycine oxidation under *in vivo* conditions, we generated an *E. coli* strain to serve as a glyoxylate biosensor (Fig. 4a). We constructed an *E. coli* strain in which the native anaplerotic reactions were deleted (Δ*ppc* Δ*pck* Δ*maeA* Δ*maeB*), such that the biosynthesis of malate from the condensation of acetyl-CoA and glyoxylate serves as the only route for the net production of TCA cycle intermediates (Fig. 4a). To enable this reaction, the gene encoding for malate synthase (glcB) was genomically overexpressed using a constitutive strong promoter (Methods). All enzymes that produce or consume glyoxylate (Δ*glcDEF* Δ*aceA* Δ*gcl* Δ*ghrA* Δ*ghrB*) were further deleted in this strain, thus insulating this compound from the rest of metabolism. The glyoxylate biosensor strain was able to grow only if glyoxylate was added to the medium (red line vs. purple line in Fig. 4b). We found that upon expression of *C. necator dadA6*, glycine could replace glyoxylate, supporting fast growth of the biosensor strain (blue line in Fig. 4b). This confirms that DadA6 can effectively catalyze glycine oxidation to glyoxylate under physiological conditions.

**Figure 4.**
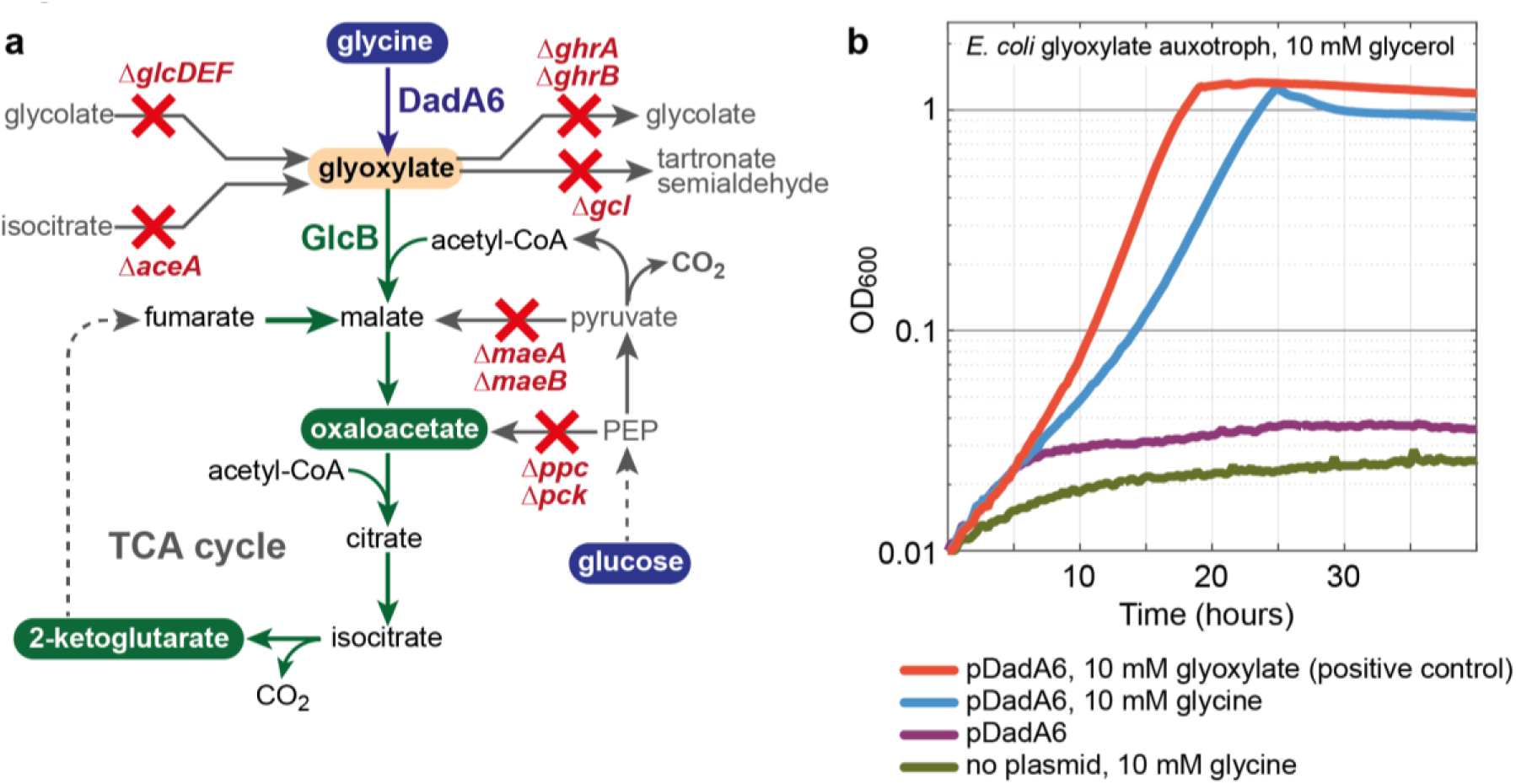
DadA6 catalyzes the *in vivo* oxidation of glycine. (**a**) An *E. coli* glyoxylate biosensor strain was constructed by deleting all anaplerotic reactions, as well as all enzymes that consumed or produced glyoxylate. Malate synthase (GlcB) was overexpres sed from the genome using a strong constitutive promoter. The growth of this strain is dependent on glyoxylate assimilation to malate to provide the essential TCA cycle intermediates oxaloacetate and 2-ketoglutarate. (**b**) Heterologous expression of *C. necator dadA6* rescued the growth of this glyoxylate auxotroph in the presence of glycine. Growth experiments were conducted in 96-well plate readers on a minimal medium (M9). Growth experiments were performed in triplicates and showed identical growth curves (±5%); hence, representative curves and average doubling times (t_D_) are shown. All experiments were repeated independently at least three times and showed highly similar growth behavior.

To further confirm that glycine assimilation occurs via its oxidation to glyoxylate and the operation of the glycerate pathway, we deleted either *dadA6* or *gcl-tsr* in the glycine-evolved strain. Either deletion completely abolished growth on glycine (red and orange lines in Fig. 3b). On the other hand, deletion of the genes encoding for the GCS, which would be essential for growth on glycine via the “serine route” (Fig 3a), did not substantially affect growth (green line in Fig. 3b). This unequivocally confirms that rather than operating the “serine route”, as in the original design of the rGlyP, *C. necator* assimilates glycine via the “glyoxylate route” (Fig. 3a).

### Growth on formate via the ‘glyoxylate’ variant of the reductive glycine pathway

We aimed to integrate the two segments of the rGlyP (Fig. 5a). We hypothesized that overexpression of the enzymes of the upper segment – converting formate to glycine – would suffice to establish growth on formate, as the accumulation of glycine would induce the glycine-assimilating segment, as shown above. First, we deleted the genes encoding for both Rubisco isozymes (*cbbSLc2* on chromosome 2 and *cbbSLp* on a megaplasmid) in a non-evolved *C. necator*^23^, thus abolishing growth on formate via the Calvin cycle. We transformed this strain with pC1 and pC2 carrying different combinations of promoters (Methods and Fig. S1, S2). After two weeks of incubation in a minimal medium with formate (10% CO_2_ and 100 mM bicarbonate) we observed growth of three cultures harboring pC1-p_14_ and pC2-p_3_. Upon reinoculation to a fresh medium with formate, these strains, which we termed CRG1 (***C***. *necator* **rG**lyP 1), were able to immediately grow on formate, albeit at a low growth rate (purple line in Fig. 5c represents one of this strains, having a doubling time 56 h). To test whether glycine assimilation proceeds via the “glyoxylate route” (Fig. 3a), as was the case when glycine was provided in the medium, we deleted *dadA6* in a CRG1 strain. This CRG1 Δ*dadA6* strain was effectively unable to grow on formate (light red line in Fig. 5c), confirming that the growth of the CRG1 strain takes place via glycine oxidation (Fig. 5a).

**Figure 5.**
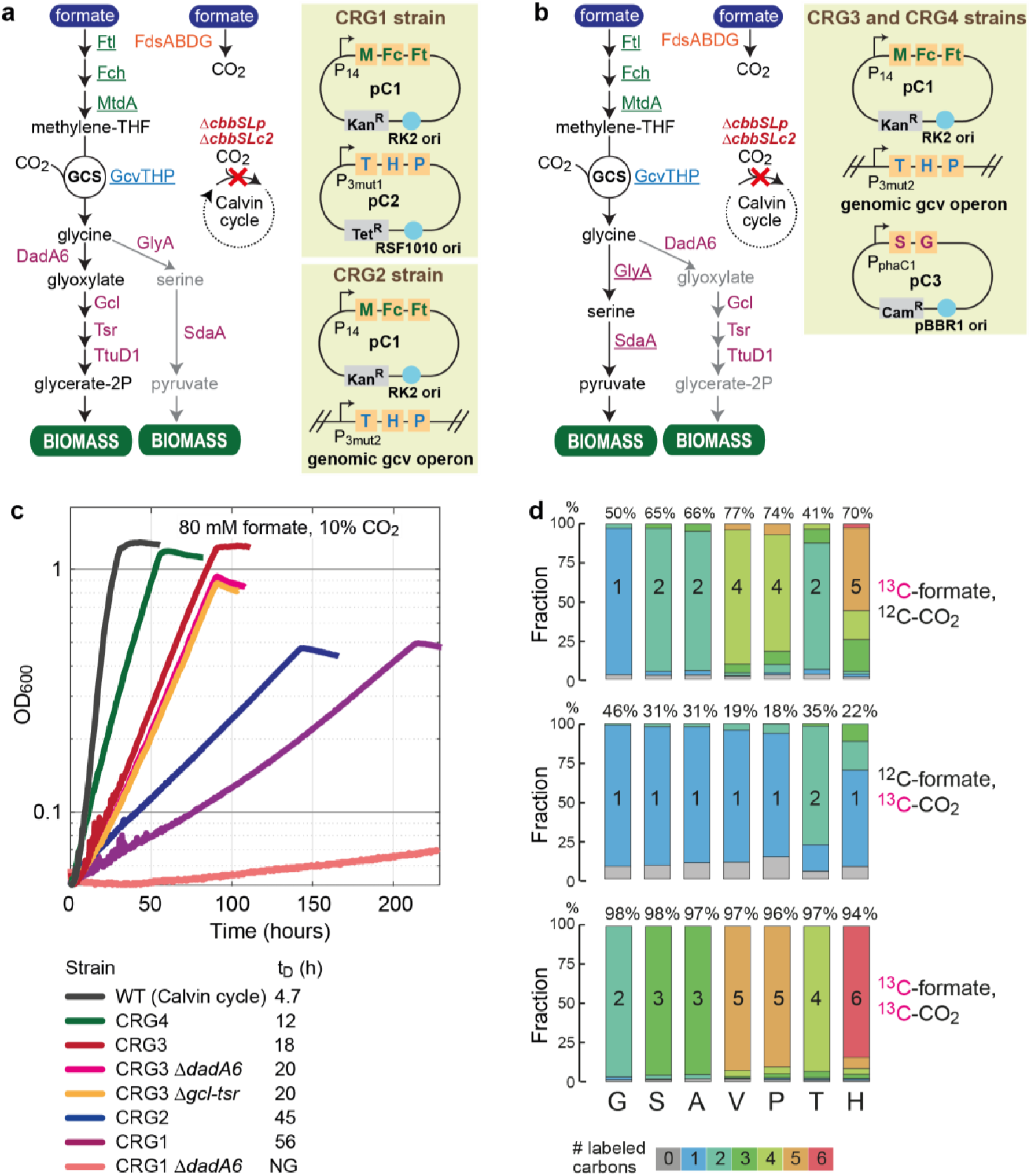
Establishing and optimizing *C. necator* formatotrophic growth via the rGlyP. (**a**) Formatotrophic growth of a *C. necator* strain (in which the Calvin cycle was disrupted) using the ‘glyoxylate’ variant of the rGly was made possible after short-term evolution upon expression of the upper segment of the pathway (underlined enzymes) either on plasmid or within the genome (‘M’ corresponds to *mtdA*; ‘Fc’ to *fch*; ‘Ft’ to *ftl*; ‘T’ to *gcvT*; ‘H’ to *gcvH*; and ‘P’ to *gcvP*). The promoter controlling the expression of the GCS gene was mutated both on a plasmid (CRG1 strain) and in the genome (CRG2 strain) (Fig. S1, S2, S3 and Supplementary Data 3). (**b**) Formatotrophic growth of a *C. necator* strain (in which the Calvin cycle was disrupted) using the ‘serine’ variant of the rGly was made possible upon expression of the lower segment of the pathway (underlined enzymes in purple) within the CRG2 strain (‘S’ to *sdaA* and ‘G’ to *glyA*). The CRG4 strain was obtained after short-term evolution of the CRG3 strain. (**c**) Growth of different strains on formate. Growth experiments were performed in 96-well plates on a minimal medium (JMM) supplemented with 80 mM formate, 100 mM bicarbonate and 10% CO_2_. Experiments were considered in triplicates that showed identical growth curves (±5%); hence, a representative curve and average doubling times (t_D_) are shown. All experiments were repeated independently at least twice and showed highly similar growth behavior. (**d**). ^13^C-labeling of proteinogenic amino acids (‘G’ corresponds to glycine; ‘S’ to serine; ‘A’ to alanine; ‘V’ to valine, ‘P’ to proline; ‘T’ to threonine; and ‘H’ to histidine) upon cultivation of the CRG4 strain on different combinations of labeled and unlabeled formate and CO_2_. Numbers written above the bars correspond to the overall fraction of labeled carbons. The labeling confirms the activity of the pathway and indicate low cyclic flux via the TCA cycle (Fig. S4). Labeling experiments are averages from duplicates.The CRG2 strain showed a faster growth on formate than the CRG1 strains, having a doubling time of 45 hours (blue line in Fig. 5c). We sequenced the genome of the CRG2 strain and identified a few mutations (Supplementary Data 4). Among these, one was directly downstream of promoter p^3^. We conducted quantitative PCR and found that the expression of *gcvT* (the first gene in the GCS operon) was an order of magnitude higher with the p_3mut2_ promoter than with the original p_3_ promoter (Fig. S3). Interestingly, the transcript level of *gcvT* in the CRG2 strain (genomic expression via p_3mut2_) was similar to that observed in the CRG1 strain when the GCS was expressed on a plasmid under the regulation of p_3mut1_ (Fig. S3).

We sequenced the CRG1 strains and found mutations both in the genome (Supplementary Data 3) and on the plasmid (Fig. S1). Several of these mutations occurred in all three CRG1 strains. One such shared mutation occurred inside the promoter p_3_ on the pC2 plasmid, resulting in the mutated promoter p_3mut1_ with an order of magnitude lower strength (Methods and Fig. S2). Another shared mutation was the deletion of *ccbR*c2, which encodes for the key activator of all Calvin cycle genes, activating both CO_2_ fixation operons on chromosome 2 and the megaplasmid^24^. The contribution of this deletion to growth might be attributed to the downregulation of phosphoribulokinase, which upon the deletion of Rubisco, generates the dead-end metabolite ribulose 1,5-bisphosphate.

As the initial high expression of the GCS seems to be deleterious (as suggested by the mutated promoter), we decided to replace its overexpression from a plasmid with genomic overexpression. We therefore cured the CRG1 strain from the pC2 plasmid and replaced the native, genomic promoter of the GCS operon with six constitutive promoters of different strength (Methods and Fig. S1, S2). We inoculated this strain in a minimal medium with formate (at 10% CO_2_ and 100 mM bicarbonate). After four weeks, we observed growth of several cultures harboring different GCS promoters. Reinoculation of these strains in fresh media with formate enabled immediate growth. The strain, in which the genomic GCS was engineered under the control of p_3_, was termed CRG2. The CRG2 strain showed the best growth and was further analyzed.

### Improved growth on formate via the ‘serine’ variant of the reductive glycine pathway

The “glyoxylate route” for glycine assimilation is less efficient than the “serine route”, as the former wastes reducing power during glycine oxidation; specifically, the expected pyruvate yield from formate using the “serine route” is >30% higher than with the “glyoxylate route” (Fig. 1b). We therefore aimed to force glycine assimilation via the “serine route”, bypassing its oxidation (Fig. 5b). We cloned the native genes encoding for serine hydroxymethyltransferase (*glyA*) and serine deaminase (*sdaA*) – the two components of the “serine route” – and assembled them into a synthetic operon on a plasmid, which we termed pC3, under the control of four different constitutive promoters of varying strength (Methods and Fig. S1, S2). We transformed the CRG2 strain with the pC3 plasmid and tested whether its growth rate was improved. We found that expression of *glyA* and *sdaA* from the medium promoter p_phaC1_ improved growth the most, decreasing doubling time to 18 hours and increasing biomass yield (i.e., final OD_600_) by more than 2-fold (red line in Fig. 5c). To check whether this strain, which we termed CRG3, is indeed independent on the “glyoxylate route” we deleted either *dadA6* or *gcl-tsr*. Neither of these deletions substantially altered the growth phenotype (pink and orange lines in Fig. 5c), confirming that the “serine route” replaced the “glyoxylate route”. The CRG3 thus assimilates formate via the rGlyP using its original, more efficient design.

To further improve growth on formate, we conducted a short-term adaptive evolution, in which, upon reaching stationary phase, the culture was reinoculated in fresh medium at OD_600_ of 0.05. After a several cycles of cultivation, the growth rate of the culture increased. We isolated a strain, termed CRG4, in which the growth rate increased by 50% (doubling time of 12 hours, green line in Fig. 5c). We sequenced the genome of strain CRG4 and found several mutations, none of which in genes or regulatory elements directly related to the rGlyP (Supplementary Data 5). We measured the exact biomass yield of the CRG4 strain and found it to be 2.6 gCDW/mol formate, similar to the biomass yield of the WT strain growing on formate via the Calvin cycle, 2.9 gCDW/mol formate.

To confirm that growth of the CRG4 strain takes place via the rGlyP we performed ^13^C-labeling experiments. We cultivated the strain with ^13^C-formate/^12^CO_2_, ^12^C-formate/^13^CO_2_, or ^13^C-formate/^13^CO_2_, and measured the labeling pattern of proteinogenic glycine, serine, alanine, valine, proline, threonine, and histidine; these amino acids either directly relate to the activity of the rGlyP or originate from different parts of central metabolism, thus providing an indication of key metabolic fluxes (Fig. S4). When cultivated on ^13^C-formate/^13^CO_2,_ all amino acids were 94-98% labeled (text above the bars in Fig. 5d), indicating that formate and CO_2_ indeed serve as the only carbon sources (as formate and CO_2_ are only 98-99% labeled, 100% labeling of the amino acids is not achievable). The different labeling patterns of glycine, serine, alanine, and valine when fed with ^13^C-formate/^12^CO_2_ or ^12^C-formate/^13^CO_2_ (Fig. 5d) match the expected pattern from the activity of the rGlyP (Fig. S4). The labeling pattern of proline and threonine, which are derived from intermediates of the tricarboxylic acid (TCA) cycle (Fig. 5d), indicate a very low cyclic flux (Fig. S4). This is consistent with the use of formate as a primary reducing power and energy source, thus making the full oxidation of acetyl-CoA unnecessary.

## Discussion

In this study, we demonstrated the successful engineering and optimization of the synthetic rGlyP in *C. necator*, replacing the Calvin cycle for supporting growth on formate. To facilitate this, we divided the pathway into two segments – (i) formate conversion to glycine and (ii) glycine assimilation to biomass – and explored the activity of each separately before combining them into a full pathway. We discovered that *C. necator* can effectively assimilate intracellular glycine into biomass via its oxidation to glyoxylate and the activity of the glycerate pathway. However, since this route is rather inefficient due to a wasteful dissipation of reducing power, we replaced it with glycine conversion to serine and pyruvate. We further demonstrated the strength of integrating both rational design and short-term evolution to optimize pathway activity. This approach enabled us to more than double the growth yield on formate and increase growth rate almost 4-fold (Fig. 5c).

During the short-term adaptive evolution, we identified several mutations that might have contributed to the improved growth. However, as manipulating *C. necator*’s genome is difficult, a systematic exploration of the contribution of each mutation to the phenotype could not be easily performed. Moreover, while shifting overexpression from a plasmid to the genome improved growth (i.e., the genes of the GCS), we were not able to replace all plasmids with genomic expression as the introduction of multiple-gene operon (e.g., *ftl-fch-mtdA*) into *C. necator*’s genome is still a challenging task. Once more effective tools for engineering the genome of this bacterium become available, it will be possible to further optimize the activity of the rGlyP and explore in detail the cellular adaptation towards efficient assimilation of formate.

Replacing *C. necator*’s Calvin cycle with the rGlyP has the potential to substantially increase biomass yield on formate. However, in this study we were able only to match the yield of the natural route. This should not come as a surprise as the bacterium is still not fully adapted to the use of the synthetic pathway. Further optimization of pathway activity, using both rational engineering and long-term evolution is expected to boost growth rate and yield of our strain.

The *C. necator* strain utilizing the rGlyP compares favorably with a recently evolved *E. coli* strain that grew on formate via the Calvin cycle with a doubling time of 18 hours^25^. Nevertheless, a recently engineered *E. coli* strain growing on formate via the rGlyP displayed faster growth (doubling time of ≈8 hours), albeit with a somewhat lower biomass yield (2.3 gCDW/mol-formate)^26^. However, in the long run – following further rational optimization and adaptive evolution – *C. necator* may outcompete *E. coli* due to a marked advantage: the use of a highly efficient, metal-dependent formate dehydrogenase (FDH). Specifically, in the *E. coli* studies, a metal-free FDH was used, which can be easily expressed in a foreign host but is limited by poor kinetics (k_cat_ ≤ 10 sec^-1^) ^27^. On the other hand, molybdenum/tungsten-dependent FDHs, as those natively used by *C. necator*^28,29^, are very fast (k_cat_ typically exceeds 100 sec^-1^), but are difficult to heterologously express to enable sufficient *in vivo* activity. At the current stage, the identity of the FDH variant might not be important, as growth is likely limited by metabolic factors other than the supply of reducing power and energy. However, as we keep improving formatotrophic growth via the rGlyP, the supply of reducing power will become more and more limiting, and the bacterium that harbors the more efficient FDH could have a clear advantage^16^.

The successful implementation of the rGlyP into both *E. coli* and *C. necator* (for which less genetic tools are available) suggests that this pathway is robust enough to be introduced to various relevant hosts. This robustness can be attributed to several factors, including the use of mostly ubiquitous enzymes, a linear structure that avoids the need for balancing fluxes within a cyclic route, and operation at the periphery of metabolism, thus negating deleterious clashes with central metabolism. The implementation of the rGlyP in multiple biotechnologically-relevant microorganisms therefore seems a viable strategy, providing flexible platforms for valorizing CO_2_-derived formate into a myriad of value-added chemicals.

## Methods

### Bacterial strains and conjugation

A *C. necator H16* strain knocked out for polyhydroxybutyrate biosynthesis (*ΔphaC1*) was used as a platform strain for engineering in this study (kindly donated by O. Lenz)^30^. *E. coli* DH5α was used for routine cloning, while *E. coli* NEB10-beta was used for cloning of larger vectors. *E. coli* S17-1 was used for conjugation of mobilizable plasmids to *C. necator* by biparental overnight spot mating. *C. necator* transconjugants were selected on LB agar plates with the appropriate selection marker and 10 µg/ml gentamycin for counter-selection of *E. coli*. A complete overview of strain genotypes used in this study can be found in Table S1.

### *C. necator* genomic gene deletions

Genomic knockouts of target genes and operons were generated using the pLO3 suicide vector (kindly donated by O. Lenz), similar to the previously described methods^31,32^. In short, homology arms upstream and downstream of the knockout site of ∼1 kb were PCR amplified by Phusion HF polymerase (Thermo Scientific). Homology arms were assembled into digested (SacI, XbaI) or PCR-amplified pLO3 backbone via Gibson Assembly (HiFi, NEB or In-fusion, Takara). *C. necator* was conjugated with the pLO3 vectors and single-cross overs were selected on tetracycline. Transconjugants were grown overnight without tetracycline to allow for the second cross-over event. These cells were plated and pure knockout clones were screened by colony PCR using OneTaq (Thermo Scientific).

### Promoter library

A constitutive promoter library was constructed using oligos for short promoter sequences and PCR amplification of longer promoter sequences. Our initial library consisted of the constitutive promoters p_2_, p_3_, p_4_ and p_14,_ which are from the p_trc_-derived library of Mutalik *et al.* developed for *E. coli*^33^. The promoters were cloned by restriction ligation into broad-host range vectors with different ori’s: pSEVA521, pSEVA531 and pSEVA551, together with a synthetic RBS and GPF (cargo #7 from SEVA system)^34^. Relative promoter strength was measured based on GFP fluorescence as explained below (Fig. S2). All four promoters expressed GFP at different strengths in *C. necator*, but we found their relative strength ranking in *C. necator* (weak to strong: p_14_→p_3_→p_4_→p_2_) to be scrambled when comparing to the previously identified order in *E. coli*^33^ (weak to strong: p_4_→p_14_→p_2_→p_3_). The promoters from this initial library appeared rather strong in *C. necator*. Furthermore, p_14_ was the weakest promoter in a vector with a RK2 origin of replication, but in vectors with higher copy number ori’s (RSF1010 and pBBR1^35^) it appeared to be a medium-strength promoter, stronger than p_3_; the other three promoters kept their respective ranking p_3_→p_4_→p_2._ (Note: *C. necator* could not be conjugated with strongest promoter p_2_ on the highest tested copy number ori pBBR1, likely due to too high expression burden). To allow better benchmarking of our library and expanding the strength range especially with weaker promoters we included in our expanded library constitutive promoters previously tested in *C. necator:* p_tac_^35–37^, p_cat_^38^, p_lac_^35–37^, p_phaC1_^35–38^ p_j5_^35^. We also included the native *C. necator* p_pgi_ promoter (phosphoglucoisomerase) as we expected that this central glycolytic promoter could be an additional interesting constitutive promoter as seen before in *E. coli* ^39^. A broader range of promoter strengths was found, including several weaker promoters, and the previously described strongest p_j5_ was also the strongest of our 10 tested promoters in the full library (Fig. S2).

### Pathway enzyme expression plasmids

Genes *ftl, fch* and *mtdA* were PCR-amplified from *M. extorquens* AM1 genomic DNA (similar GC-content to *C. necator*, so no codon optimization needed). A synthetic RBS was included for each gene by PCR, which was designed by RBS Calculator^40^ to have a medium-strength of ∼30,000 arbitrary units, taking into account the context of the 5’ UTR and start of the gene. The genes were assembled using Gibson assembly in pSEVA221 vectors with four different promoters (p_2_, p_3_, p_4_ and p_14_) generating a library of pC1 vectors. Genes *gcvT* and *gcvHP* were PCR amplified from the *C. necator* genome and synthetic RBSs were included for *gcvT* and *gcvH* designed as described above. The synthetic operon was assembled via Gibson assembly in pSEVA551 using three different promoters (p_2_, p_3_, p_4_) generating the pC2 vectors. pC3 was constructed by PCR amplication of *C. necator’s sdaA* and *glyA* with their native RBSs and assembled in pSEVA331 with the promoters p_3_, p_4_, p_cat_, p_phaC1_, the latter two weaker promoters were included as our previous experience with pC1 and pC2 showed that the weaker promoters of the library gave better growth phenotypes. pDadA6 was generated for heterologous DadA6 expression in an *E. coli* glyoxylate biosensor, the *C. necator dadA6* gene was amplified with a synthetic RBS and assembled into pSEVA531 with the strong *E. coli* promoter p_3_.

### *gcvTHP* operon promoter exchange

To optimize overexpression of the native *gcvTHP* operon from the genome the native promoter was exchanged by 6 different promoters ranging from weak to strong (p_cat_, p_phaC1_, p_3_, p_4_, p_2_, p_j5_) as we had no good indication of what level of genomic expression was desired. To allow for promoter exchange, we constructed a pLO3 suicide vector with ∼1000 bp homology arms flanking the native p_gcvTHP_ promoter. In between the homology arms different PCR-amplified promoters were inserted by restriction digestion (using AscI/XbaI). The knock-in protocol was the same as for the knockouts in *C. necator* described above.

### *E. coli* glyoxylate biosensor strain construction

The *E. coli* SIJ488 strain based upon K-12 MG1655^41^, was used for the generation of the glyoxylate biosensor strain. SIJ488 is engineered to carry the gene deletion machinery in its genome (inducible recombinase and flippase). All gene deletions were carried out by successive rounds of λ-Red recombineering using kanamycin cassettes (FRT-PGK-gb2-neo-FRT (KAN), Gene Bridges, Germany) or chloramphenicol cassettes (pKD3^42^) as described before^43^. Homologous extensions (50 bp) for the deletion cassettes were generated by PCR. The sensor strain required overexpression of the malate synthase (*glcB*), hence, the endogenous gene was amplified from *E. coli* genomic DNA using a two-step PCR (to remove cloning system relevant restriction sites^39^ – in this case a single site). The *glcB* gene was subsequently cloned into cloning vector pNivB^44^ using restriction and ligation (Mph1103I/XhoI), generating pNivB-glcB. The *glcB* gene was subsequently cloned from pNivB-*glcB* into a BioBrick adapted pKI^39^ suicide vector for integration at the Safe Site 9^45^ under the control of a strong constitutive promoter in the *E. coli* genome using enzymes *Eco*RI and *Pst*I, resulting in pKI-SS9-B-glcB. In brief, the knock-in system relies on conjugation of the suicide vector via a ST18 *E. coli* strain which requires 5-aminolevulinic acid for growth^46^ and a sucrose (*sacB*) counterselection system (system described in full in^39^).

### Growth medium and conditions

*C. necator* and *E. coli* were cultivated for routine cultivation and genetic modifications on Lysogeny Broth (LB) (1% NaCl, 0.5% yeast extract and 1% tryptone). When appropriate the antibiotics kanamycin (100 µg/mL for *C. necator* or 50 µg/mL for *E. coli)*, tetracycline (10 µg/mL), chloramphenicol (30 µg/mL), ampicillin (100 µg/mL for *E. coli*) or gentamycin (10 µg/mL for *C. necator*) were added. Routine cultivation was performed in 3 mL medium in 12 mL glass tubes in a shaker incubator at 240 rpm. *C. necator* was cultivated at 30°C and *E. coli* at 37°C.

Growth characterization experiments of *C. necator* were performed in J Minimal Medium (JMM) a medium previously optimized for formatotrophic growth^12^. For formatotrophic growth 80 mM sodium formate was added, as well as 100 mM sodium bicarbonate, pH was adjusted to 7.2 and incubation was performed under a headspace of 10% CO_2_. No antibiotics were added during growth characterization experiments. Growth of the glycine auxotroph was supplemented with 8 mM glycine when appropriate, while the glycine evolved strain was grown with 40 mM glycine. Organic carbon sources fructose or pyruvate were used for pre-cultures or control cultures at 20 and 40 mM respectively.

The *E. coli* glyoxylate biosensor was grown on M9 minimal medium (50 mM Na_2_HPO_4_, 20 mM KH_2_PO_4_, 1 mM NaCl, 20 mM NH_4_Cl, 2 mM MgSO_4_ and 100 μM CaCl_2_) with trace elements (134 μM EDTA, 13 μM FeCl_3_·6H_2_O, 6.2 μM ZnCl_2_, 0.76 μM CuCl_2_·2H_2_O, 0.42 μM CoCl_2_·2H_2_O, 1.62 μM H_3_BO_3_, 0.081 μM MnCl_2_·4H2O) with glycerol (10 mM) and supplemented with 10 mM glycine or glyoxylate when appropriate.

Growth curves were monitored during growth in 96 well-plates (Nunc transparent flat-bottom, Thermo Scientific). Precultured strains were washed twice and inoculated at an OD_600_ of 0.01 (0.05 for growth on only formate). Each well contained 150 μl of culture topped with 50 μl of transparent mineral oil (Sigma-Aldrich) to avoid evaporation (O_2_ and CO_2_ can freely diffuse through this layer). Plates were incubated with continuous shaking (alternating between 1 minute orbital and 1 minute linear) in a BioTek Epoch 2 plate reader. OD_600_ values were measured every 12 minutes. Growth data were processed by a Matlab script, converting plate reader OD_600_ to cuvette OD_600_ by multiplication by a factor 4.35. All growth experiments were performed in at least in triplicates, and the growth curves shown are representative curves.

### Whole-genome sequencing

*C. necator* genomes (and plasmids) were extracted for whole-genome sequencing using the GeneJET Genomic DNA Purification Kit (Thermo Fisher). Samples were sent for library preparation (Nextera, Ilumina) and sequencing at an Ilumina NovoSeq 6000 platform to obtain 150bp paired-end reads (Baseclear, Netherlands or Novogene, UK). Samples were paired, trimmed and assembled to the *C. necator* reference genome using Geneious 8.1 software. Mutations (frequency above >60%) were identified based on comparative analysis with the parental strains.

### Transcriptomic sequencing

Cells were grown in triplicates on JMM 40 mM glycine and 40 mM pyruvate till mid-log phase and 1-2 mL of cultures were harvested and stabilized directly by RNA Protect Bacteria Kit (Qiagen). Cells were then lysed using lysozyme and beating with glass beads in a Retschmill (MM200) for 5 minutes at 30 hertz. RNA was purified using the RNeasy Mini kit (Qiagen) following manufacturer’s protocol including on-column DNAase digestion (DNase kit Qiagen). rRNA depletion (RiboZero kit), cDNA library preparation, and paired-end 150 bp read sequencing (Illumina HiSeq 3000) was performed by the Max Planck Genome Centre Cologne, Germany. Sequence data of all samples were mapped with STAR v2.5.4b ^47^ using default parameters. Ensembl version 38 genome reference in FASTA format and Ensembl version 38 cDNA Annotation in GTF format were used for genome indexing with adapted parameters for genome size (--genomeSAindexNbases 10) and read length (--sjdbOverhang 150) to ReadsPerGene files.

### Promoter characterization with GFP

*C. necator* strains overexpressing GFP and control cells without a plasmid were grown at least in triplicates in JMM with 20 mM fructose in 96-well plates as describe above. OD_600_ and fluorescence (excitation 485 nm, emission 512 nm, gain 70) were detected in a Tecan Infinite 200 Pro plate reader. The strength of promoters was assessed based on fluorescence normalized per cell density (RFU/OD_600_) after 100 hours of cultivation, when all cultures were stationary. Values were corrected for the background fluorescence of cells without GFP.

### Promoter characterization by quantitative PCR

Cells with plasmid or genomic *gcv* from different promoters expression were grown at least in triplicates on JMM 20 mM fructose till mid-log phase and RNA isolation was performed as explained above (transcriptome sequencing). cDNA synthesis was performed using the Quantabio qScript cDNA Synthesis Kit according to the manufacturer’s protocol. qPCR was performed with primers for *gcvT (5’-*caggacatggatatcaacacctc and 5’-aagtcacgctcgctctgc, product 77 bp) using the previously published primers for *gyrB* (gyrase) as a control gene to normalize expression levels^48^. Quantitative PCR was performed using SYBR Green /ROX qPCR Master Mix (Thermo Scientific) in an Applied Biosystems 7900HT Fast Real-Time PCR System according to the manufacturer’s instructions. The qPCR protocol was: 10 min at 50°C, 5 min at 95°C, 40 cycles of 10 s at 95°C and 30 s at 60°C, and finally 1 min at 95°C. A dilution series of cDNA was made to generate a standard curve to correct for PCR efficiency. Data were analyzed as described in literature^49^.

### Labeling of proteinogenic amino acids

For labeling analysis of glycine auxotroph strains and CRG4, precultures were performed in JMM with 80 mM formate (and 20 mM fructose for glycine auxotroph) and 10% CO_2_ in the headspace. Cells were then washed twice and re-inoculated at an OD_600_ of 0.01 into the same media as the precultures, but when applicable sodium formate or CO_2_ were replaced by ^13^C sodium-formate (Sigma-Aldrich) and/or ^13^CO_2_ (Cambridge Isotope Laboratories). Cells were incubated in 3 mL in tubes, and for cultures with ^13^CO_2_ tubes were placed in an airtight 6L-dessicator (Lab Companion) filled with 10% ^13^CO_2_ and 90% air on a shaker platform (180 rpm). When reaching stationary phase 1 mL of culture was harvested, washed twice with dH_2_O and resuspended in 1 mL 6 M HCl. Cells were hydrolyzed overnight at 95°C, and then evaporated under an airstream for 2-4 hours after which the hydrolysate was resuspended in 1 mL dH_2_O. The hydrolysate was analyzed using ultra-performance liquid chromatography (UPLC) (Acquity, Waters) using a HSS T3 C18-reversed-phase column (Waters). The mobile phases were 0.1% formic acid in H_2_O (A) and 0.1% formic acid in acetonitrile (B). The flow rate was 400 µL/min and the following gradient was used: 0-1 min 99% A; 1-5 min gradient from 99% A to 82%; 5-6 min gradient from 82% A to 1% A; 6-8 min 1% A; 8-8.5 min gradient to 99% A; 8.5-11 min – re-equilibrate with 99% A. Mass spectra were acquired using an Exactive mass spectrometer (MS) (Thermo Scientific) in positive ionization mode. Data analysis was performed using Xcalibur (Thermo Scientific). The identification amino acids was based on retention times and m/z values obtained from amino acid standards (Sigma-Aldrich).

### Dry weight yield experiments

WT and CRG4 cells were cultured in duplicates in 30 mL JMM medium with 80 mM formate with 100 mM sodium bicarbonate and 10% CO_2_ in 125 mL Erlenmeyer flasks. When cells were close to stationary phase the cultures were harvested and washed twice with dH_2_O and finally resuspended in 2 mL dH_2_O. Washed cell suspension were added to pre-weighed aluminum cups. Cells were dried for 24 hours at 90°C and cell weight was determined by weighing again. Residual concentrations of formate were checked by a Formate Assay Kit following the manufacturer’s protocol (Sigma-Aldrich).

## Acknowledgements

We thank Änne Michaelis for help with the LC-MS experiments. Steffen N. Lindner helped with the engineering of the *E. coli* glyoxylate biosensor strain. We thank Victor de Lorenzo’s lab for providing us with the SEVA vectors and Tobias Erb for providing *Methylobacterium extorquens* genomic DNA. We thank the Max Planck Genome Centre Cologne (http://mpgc.mpipz.mpg.de/home/) for performing the RNA-sequencing in this study. Axel Fisher assisted with the bioinformatic analysis of the genome and transcriptome data. Eleftheria Saplaoura and Selcuk Aslan assisted with the quantitative PCR experiments. We thank Irene Sánchez-Andrea and Beau Dronsella for critical reading of this manuscript. This project was funded by the Max Planck Society. N.J.C. is funded by the Dutch Organization of Science (NWO) by a Rubicon (019.163LW.035) and a Veni (VI.Veni.192.156) fellowship.

## Author contributions

This study was designed and supervised by N.J.C. and A.B.-E. Experiments were performed by N.C.J., G.B.-F, C.A.R.C., M.F.-B., L.F., N.G.-L., A.D.M., M.M.-P., W.N., G.S., J.V., S.dV. and S.Y. Data analysis was performed by all authors. The manuscript was written by N.J.C. and A.B.-E.

## Data availability

Raw genome sequencing and transcriptome data will be deposited at NCBI. Complete information on the experimental setup as well as detailed results are available from the corresponding author upon reasonable request.

## Supplementary Figures

**Supplementary Figure 1.**
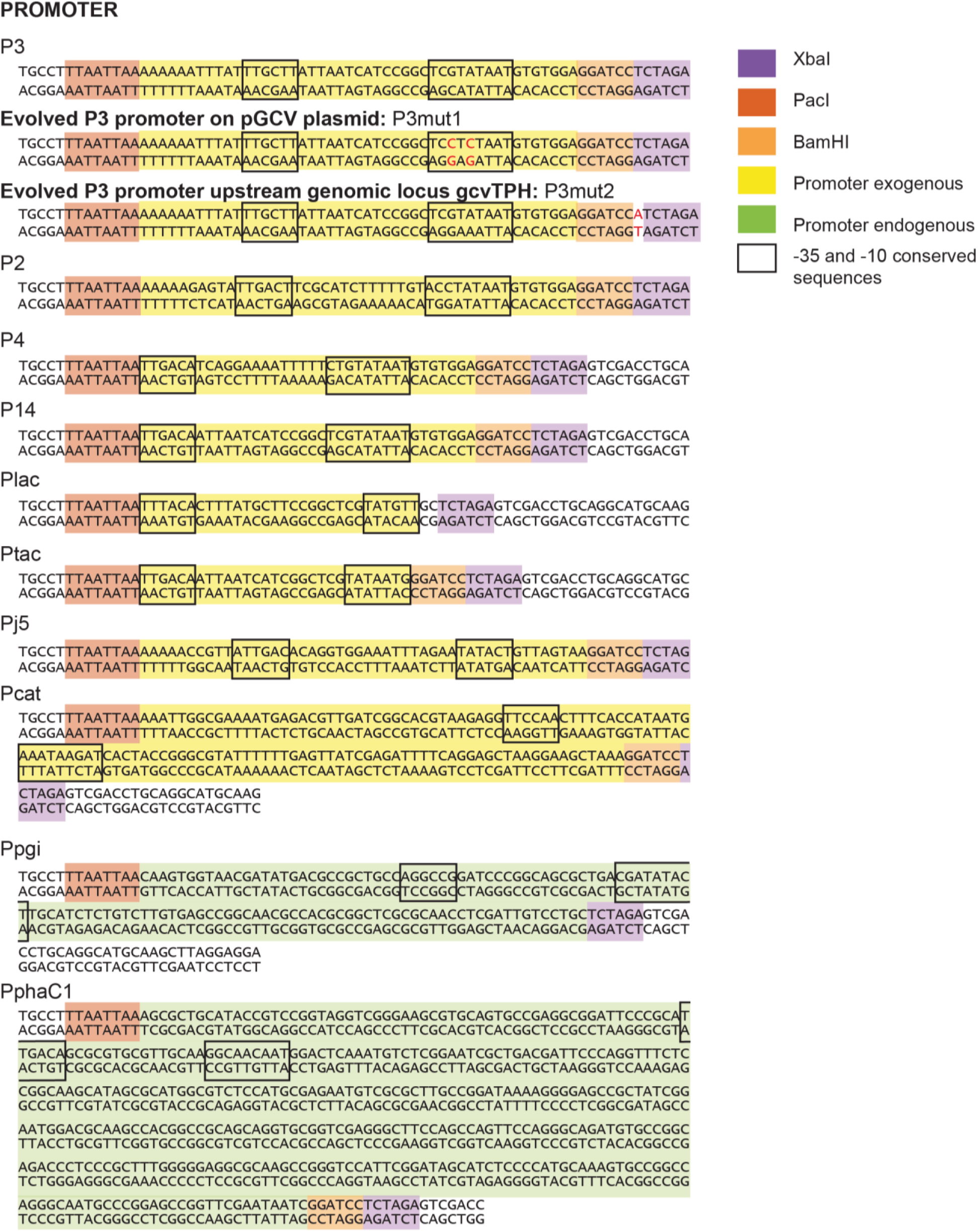
Constitutive promoters used and evolved during this study. Sequences of a set of constitutive *E. coli* promoters (p_2_, p_3_, p_4_ and p_14_), the *C. necator* p_pgi_ (phosphoglucoisomerase) promoter, and several constitutive promoters previously demonstrated to work in *C. necator*. During selection for growth on formate, promoter p_3_ was mutated to p_3mut1_ on the plasmid and to p_3mut2_ on the genome (mutations are in red).

**Supplementary Figure 2.**
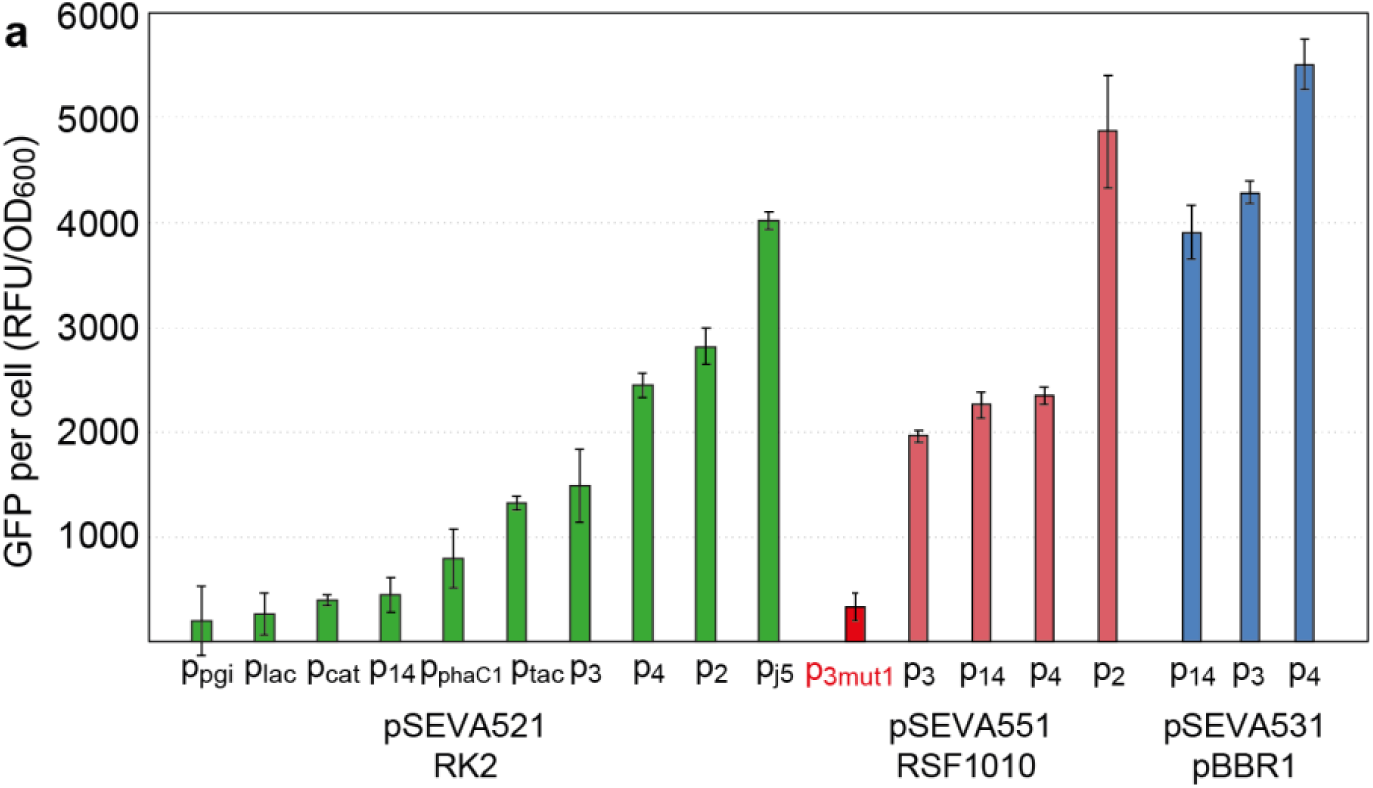
Strength of constitutive promoters on different plasmids. Promoter sequences are shown in Supplementary Figure 1. p_3mut1_ is a mutated version of p_3_ which emerged during selection for growth on formate. GFP was used as a reporter. GFP expression was measured after 100 hours growth on a minimal medium (JMM) with 20 mM fructose in 96-well plates. Experiments were conducted in triplicates. Fluorescence was normalized to OD_600_ and to auto-fluorescence of a wild-type *C. necator*.

**Supplementary Figure 3.**
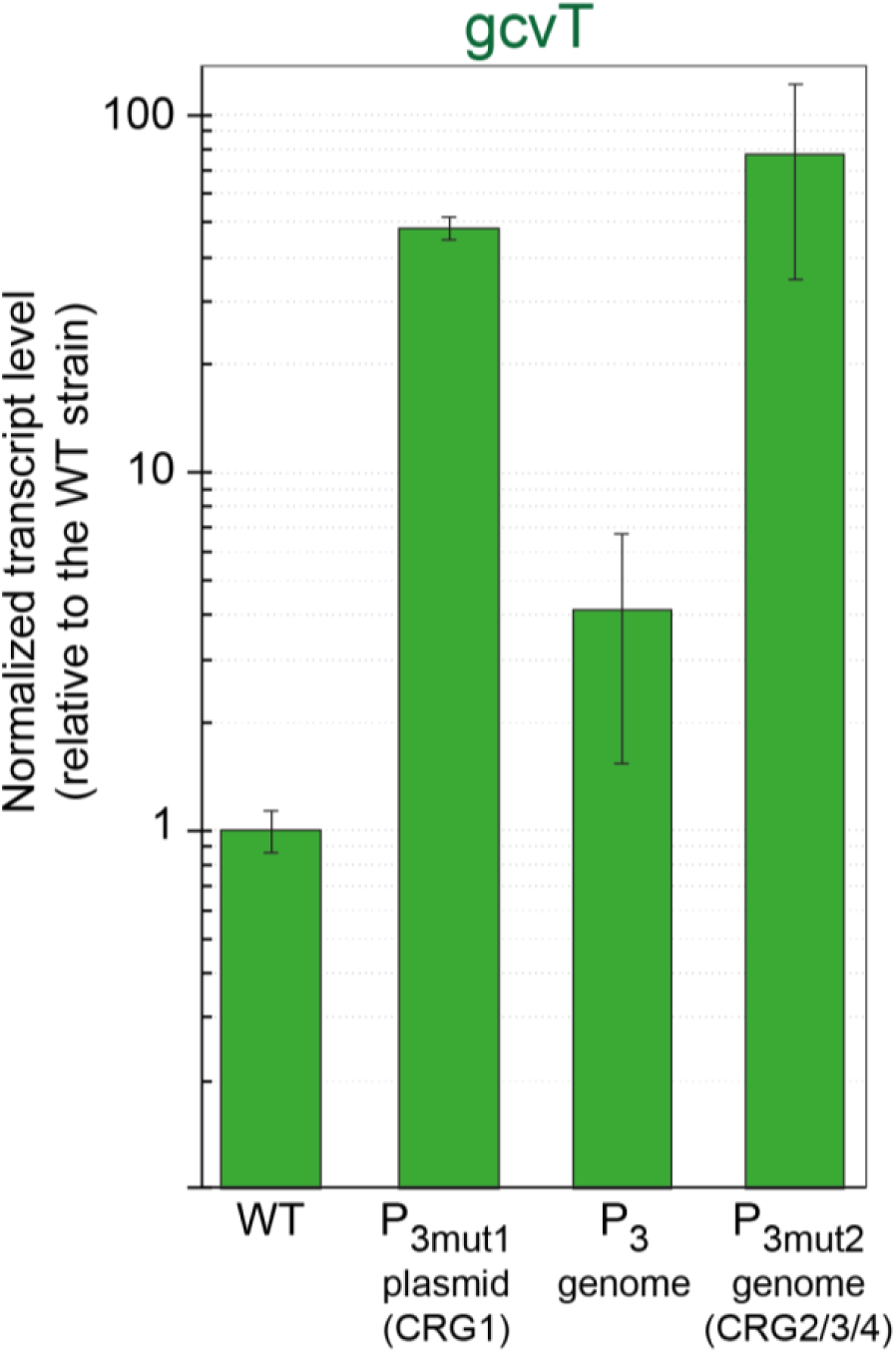
Quantitative PCR reveals transcription strength of gcvT from different p_3_ and its mutated variants. Experiments performed in triplicates and normalized according to the housekeeping gene *gyrA*. The results shown are normalized the *gcvT* transcript level in a wild-type *C. necator*.

**Supplementary Figure 4.**
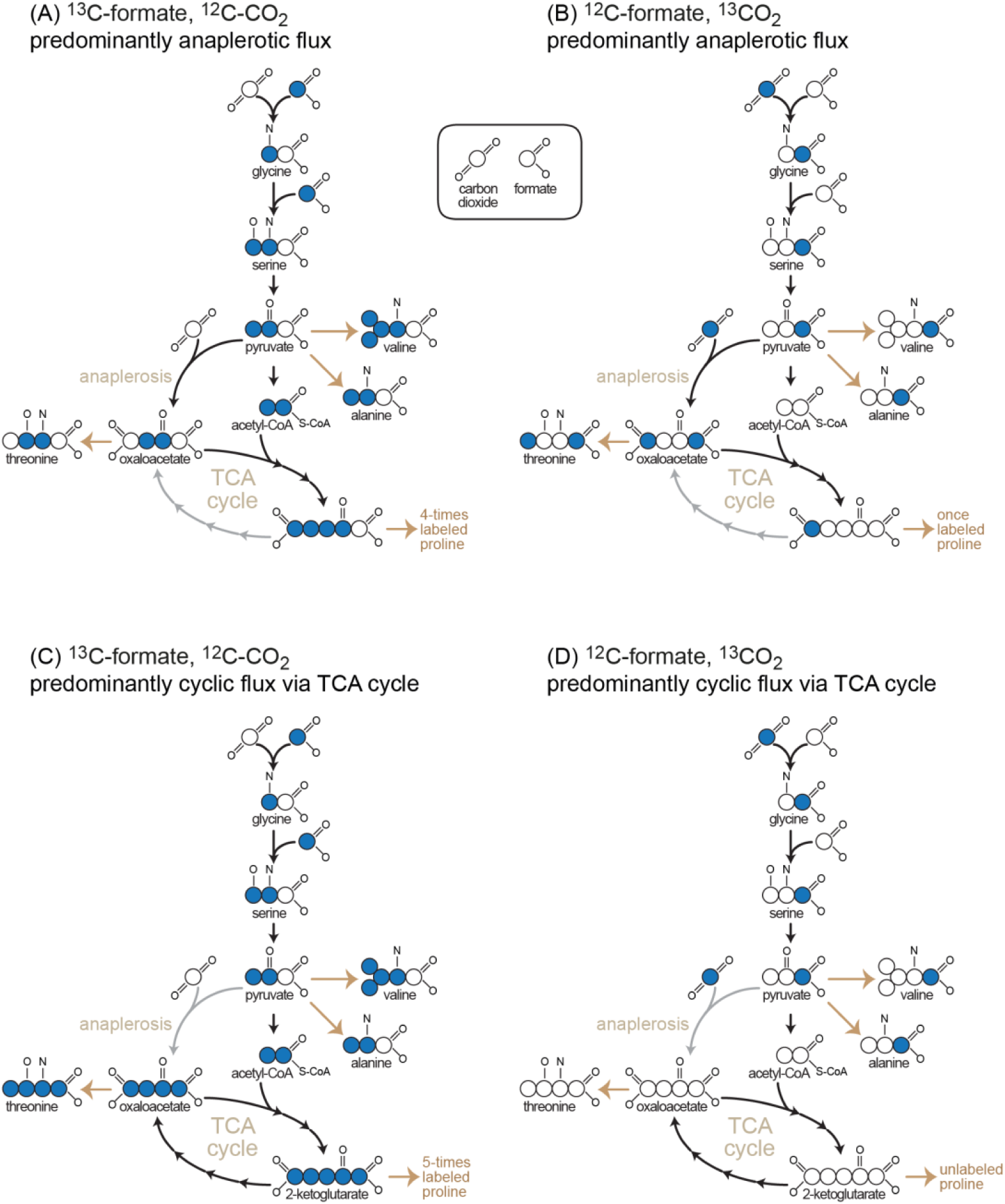
Expected labeling of proteinogenic amino acids upon feeding ^13^C-formate/^12^C-CO_2_ or ^12^C-formate/^13^C-CO_2_ according to different metabolic scenarios.

## Supplementary Tables

**Supplementary Table S1:**
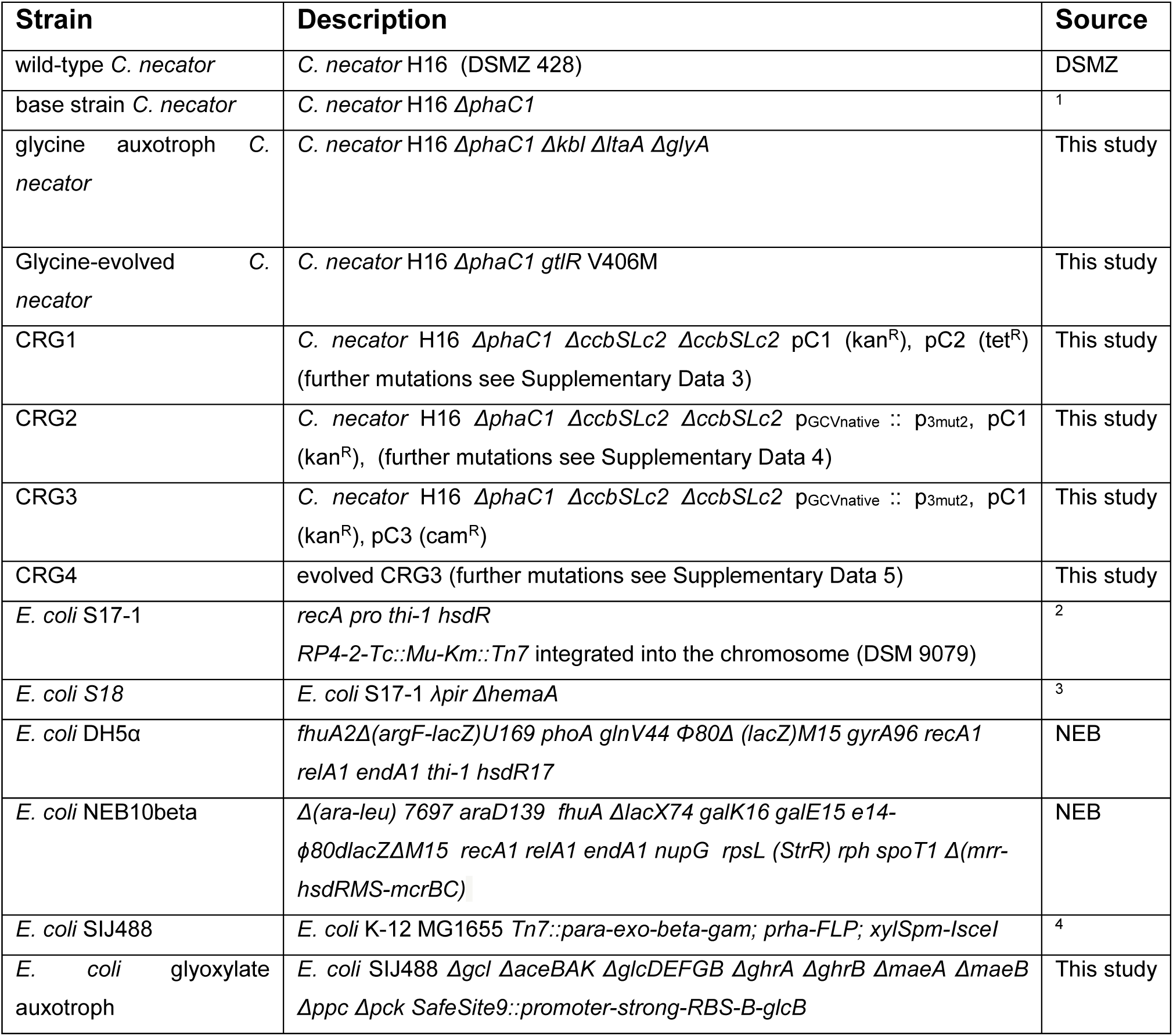
Strains used in this study.

**Supplementary Table S2:** Plasmids used in this study.

| Plasmid | Description | Source |
| --- | --- | --- |
| pLO3 | Suicide vector for knock-outs and promoter knock-ins in *C. necator* with multiple cloning site to clone homology arms, tetracycline resistance, pMB1 replication origin, RP4 origin of transfer and, *B. subtilis* *sacB* counter-selection marker | 5 |
| pSEVA221 | Standardized vector with multiple cloning site, kanamycin resistance, RK2 replication origin, and RP4 origin of transfer | 6 |
| pSEVA331 | Standardized vector with multiple cloning site, chloroamphenicol resistance, pBBR1 replication origin, and RP4 origin of transfer | 6 |
| pSEVA521 | Standardized vector with multiple cloning site, tetracycline resistance, RK2 replication origin, and RP4 origin of transfer | 6 |
| pSEVA521 | Standardized vector with multiple cloning site, tetracycline resistance, pBBR1 replication origin, and RP4 origin of transfer | 6 |
| pSEVA551 | Standardized vector with multiple cloning site, tetracycline resistance, RSF1010 replication origin, and RP4 origin of transfer | 6 |
| pSEVA637 | Vector with GFP, gentamycin resistance, pBBR1 replication origin, and RP4 origin of transfer | 6 |
| pSEVA521-GFP | pSEVA521 backbone with varying promoters, synthetic *C. necator* RBS and GFP | This study |
| pSEVA551-GFP | pSEVA551 backbone with varying promoters, synthetic *C. necator* RBS and GFP | This study |
| pC1 | pSEVA221 backbone with varying promoters (P<sub>14</sub>, P<sub>3</sub>, P<sub>4</sub>, P<sub>2</sub>) and synthetic RBSs (RBS Calculator ~30,000 au) with *M. extorquens mtdA*, *fch* and *ftl* | This study |
| pC2 | pSEVA551 backbone with varying promoters (P<sub>3mut1</sub>, P<sub>3</sub>, P<sub>4</sub>, P<sub>2</sub>) and synthetic RBSs (RBS Calculator ~30,000 au) upstream of *C. necator gcvT*, *gcvH* and native RBS + *gcvP* | This study |
| pC3 | pSEVA551 backbone with varying promoters (P<sub>cat</sub>, P<sub>Phac</sub>, P<sub>3</sub>, P<sub>4</sub>) and native RBSs with *C. necator sdaA* and *glyA* | This study |
| pDadA6 | pSEVA531 backbone with P<sub>3</sub> promoter and native RBS and *C. necator dadA6* | This study |
| pNivB | Cloning vector for BioBrick system with RBS-B, ampicilin resistance | 7 |
| pNivB-glcB | pNivB with *E. coli glcB* gene | This study |
| pKI | Conjugation/suicide vector for knockins, chloroamphenicol resistance, RK6 replication origin, *B. subtilis sacB* counter-selection marker, | 8 |
| pKI-SS9-B-glcB | pKI with homology arms (~600 bp) for safe-spot 9 integration, integration cassette with promoter S, RBS-B and *E. coli glcB* gene | This study |

